# Microbiota and stress: a loop that impacts memory

**DOI:** 10.1101/2021.06.04.447027

**Authors:** Narjis Kraimi, Flore Lormant, Ludovic Calandreau, Florent Kempf, Olivier Zemb, Julie Lemarchand, Paul Constantin, Céline Parias, Karine Germain, Christèle Dupont, Sylvie Rabot, Catherine Philippe, Aline Foury, Marie-Pierre Moisan, Anaïs Vitorino Carvalho, Vincent Coustham, Hugues Dardente, Philippe Velge, Patrice Cousin, Thierry Chaumeil, Christine Leterrier

## Abstract

Chronic stress and the gut microbiota appear to comprise a feed-forward loop, which contributes to the development of depressive disorders. Evidence suggests that memory can also be impaired by either chronic stress or microbiota imbalance. However, it remains to be established whether these could be a part of an integrated loop model and be responsible for memory impairments. To shed light on this, we used a two-pronged approach in Japanese quail: first stress-induced alterations in gut microbiota were characterized, then we tested whether this altered microbiota could affect brain and memory function when transferred to a germ-free host. The cecal microbiota of chronically stressed quails was found to be significantly different from that of unstressed individuals with lower α and β diversities and increased *Bacteroidetes* abundance largely represented by the *Alistipes* genus, a well-known stress target in rodents and humans. The transfer of this altered microbiota into germ-free quails decreased their spatial and cue-based memory abilities as previously demonstrated in the stressed donors. The recipients also displayed increased anxiety-like behavior, reduced basal plasma corticosterone levels and differential gene expression in the brain. Furthermore, cecal microbiota transfer from a chronically stressed individual was sufficient to mimic the adverse impact of chronic stress on memory in recipient hosts and this action may be related to the *Alistipes* genus. Our results provide evidence of a feed-forward loop system linking the microbiota-gut-brain axis to stress and memory function and suggest that maintaining a healthy microbiota could help alleviate memory impairments linked to chronic stress.

## Introduction

An emerging body of literature recognizes the microbiota-gut-brain axis (MGBA) as a complex and bidirectional network of interactions between the gut microbiota and the brain impacting brain health and cognitive function ^1, 2^. In the last decade, several human and animal studies have reported links between the gut microbiome and brain-related diseases like anxiety, depression or memory deficits by demonstrating major effects of gut microbiota manipulation on these disorders ^3–7^.

Whether modifications in microbiota composition impact brain function remains elusive although the involvement of immune (proinflammatory cytokines), neural (spinal and vagus nerves), metabolic (short-chain fatty acids), endocrine and neurotransmitter pathways have A recent study ^8^ provided mechanistic evidence suggesting that stress, diet and gut microbiota generate a pathological feedforward loop that contributes to depressive disorders *via* the central endocannabinoid system. However, such a loop has not been demonstrated for other disorders that can be induced by stress, for instance cognitive alterations. Indeed, chronic stress is known to induce dysbiosis in the gut characterized by changes in gastrointestinal motility and increased intestinal permeability leading to a “leaky gut” allowing bacteria and pathogens to cross the epithelial barrier. In most cases, these changes alter gut bacterial composition modifying abundances of *Firmicutes* and *Bacteroidetes* species including a decrease in *Lactobacillus* and *Porphyromonadaceae* or an increase in *Clostridium* and *Oscillibacter* ^4, 9^.

Chronic stress has also well-known negative effects on memory ^10–15^. Gut microbiota may contribute to these effects of chronic stress on memory ^16^. For example, Li et *al*. (2009) showed an improvement in spatial memory abilities, measured using the hole-board apparatus, in mice with dietary-induced shifts in bacteria diversity ^17^. A high-fat diet also led to alterations in gut microbiota composition and memory impairments in mice subjected to the Morris water maze test or the fear conditioning test ^18, 19^. In addition, comparisons between specific pathogen-free mice and germ-free mice significantly helped to highlight the link between the MGBA and memory. Gareau et *al*. ^20^ demonstrated in 2011, a lack of memory in germ-free mice in the T-maze test and novel object test in situations with or without stress. In 2018, Lu and his colleagues ^21^ also showed significant deficits of memory in germ-free mice, which supports the important role of the microbiota in memory development. More recently an inoculation of germ-free mice with *Lactobacillus* species has also been suggested to improve short-term memory in the passive avoidance memory test ^22^. Indeed, many studies have provided evidence of the positive effects of *Lactobacillus* and *Bifidobacterium* probiotic supplementation on memory capacities in mice using the Y-maze and Barnes maze tests, object recognition test, or fear conditioning test ^23–25^, but also in rats in the Morris water maze and object recognition tests ^26–28^ and in human volunteers with several memory questionnaires ^29^. The interplay between gut microbiota and memory has also been demonstrated in the nonalcoholic fatty liver disease which is characterized by hepatic fat accumulation and is associated with central obesity and diabetes since probiotics can mitigate disturbances in spatial working memory and animal recognition that are encountered in such a syndrome ^30^. Conversely, administration of antibiotics induces deleterious effects on memory as shown in mice subjected to the social transmission of food preference test and novel object recognition test ^1, 31^.

Although the links between chronic stress and memory and gut microbiota and memory have been demonstrated, the question still remains as to whether or not chronic stress could induce memory impairments *via* gut microbiota changes alone. The aim of the present study was to provide evidence of a feed-forward loop system linking the MGBA to stress and memory function showing that a chronic stress state induces gut dysbiosis which in turn may affect the brain and memory function. Japanese quails were used because we have already shown that their anxiety-like behavior and memory properties are impacted by a chronic stress procedure (unpredictable repeated negative stimuli for 21 days) ^32, 33^ and gut microbiota manipulations (germ-free model, microbiota transfer and probiotic supplementation) ^34–37^. Moreover, Japanese quail have recently been suggested as a relevant model to study the involvement of gut microbiota in stress processes ^38^. Here, the approach of cecal microbiota transfer (CMT) was used involving the transfer of microbiota from a chronically-stressed individual to germ-free naïve quails to investigate whether the CMT induced any negative consequences on quails’ spatial and cue-based memory abilities as previously demonstrated in the stressed donors ^32^. Additional analysis of plasma corticosterone levels, short-chain fatty acid activity, KEGG (Kyoto Encyclopedia of Genes and Genomes) pathway predictions of microbiome and gene expression in the brain were carried out to reveal a stress loop linking the gut microbiota to memory function.

## Results

### Stress-induced alterations in gut microbiota composition are transferred via cecal microbiota transfer

We first analysed the effects of the chronic stress procedure on the composition of quails’ cecal microbiota. A total of 415 OTUs were found among the samples. Higher alpha diversities were observed in the unstressed quails (Shannon index: 3.54 ± 0.12 and 3.11 ± 0.07 for the unstressed and stressed quails respectively; Inverse Simpson index: 14.23 ± 2.38 and 6.74 ± 0.52 for the unstressed and stressed quails respectively, **Figure 1a**). Furthermore, differential abundance assessed at the phylum level revealed higher relative abundances of the *Firmicutes* in the unstressed quails (*p* < 0.01) and of the *Bacteroidetes* in the stressed quails (*p* < 0.05) (**Figure 2a**). At the genus level, differential abundance was observed for only one genus, *Alistipes* sp. (*p* < 0.05; **Figure 2b**), out of the 69 observed in our dataset. This genus was mainly represented by the OTU1 (85.5% of the sequences assigned to *Alistipes* sp. with > 99% identity) found to be more abundant in the stressed quails (relative abundances: 35.2% ± 2.0% and 0.7% ± 0.5% respectively in the stressed and unstressed quails).

**Figure 1:**
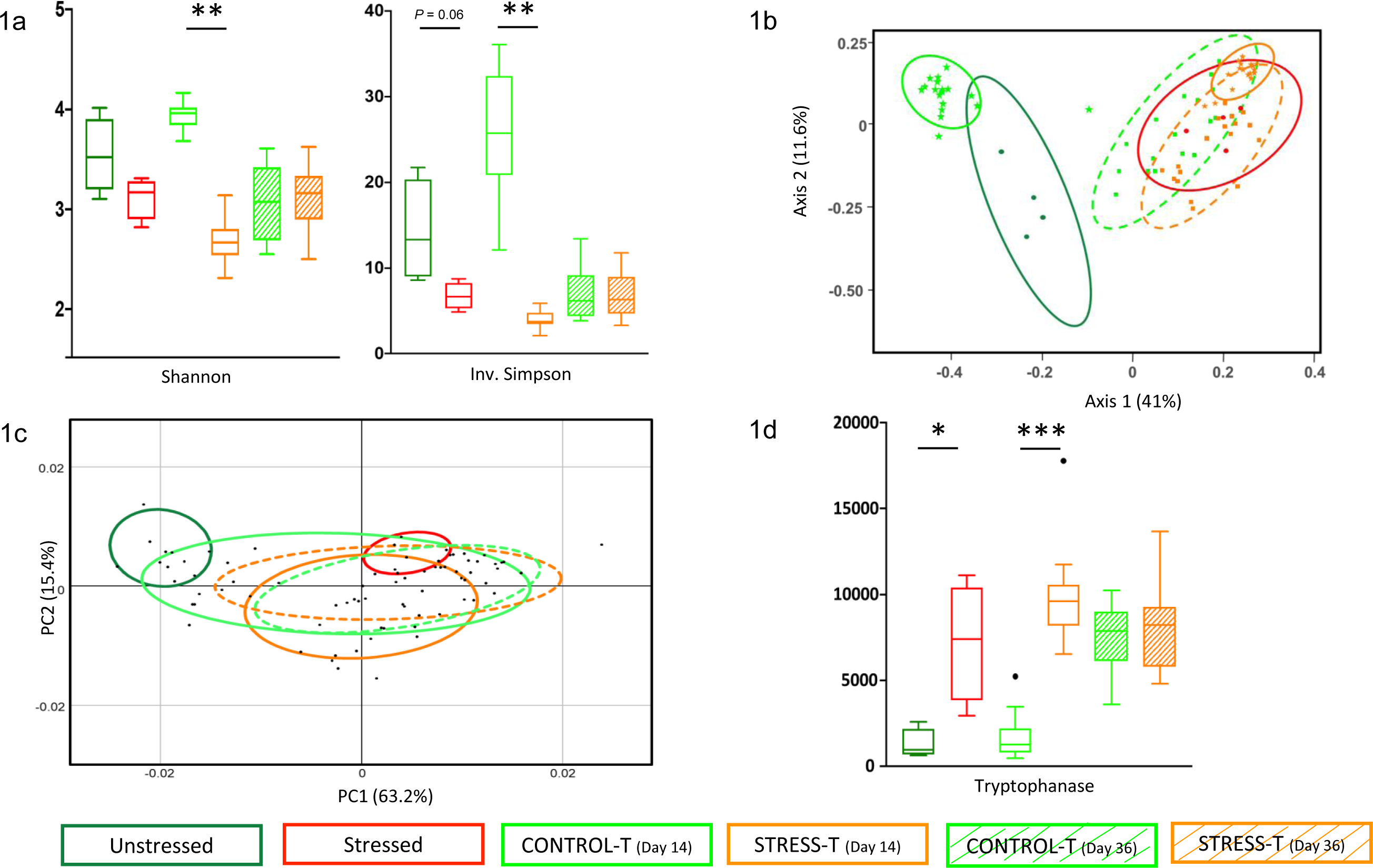
Comparison of the operational taxonomic units in cecal contents of Unstressed (n = 4) and Stressed (n = 4), CONTROL-T (n = 18) and STRESS-T (n = 17) groups at Day 14 and CONTROL-T (n = 16) and STRESS-T (n = 18) groups at Day 36 for (**a**) Shannon and Inverse Simpson α-diversity indexes, (**b**) non-Metric Multidimensional Scaling (NMDS) representation of Bray–Curtis distances, (**c**) PCA visualization of the pathway enrichment analysis derived from the taxonomic profiles, (**d**) Function prediction levels of the tryptophanase activity (EC:4.1.99.1). The results are expressed as mean values ± SEM. * *p* < 0.05, ** *p* ≤ 0.01, *** *p* ≤ 0.0001.

**Figure 2:**
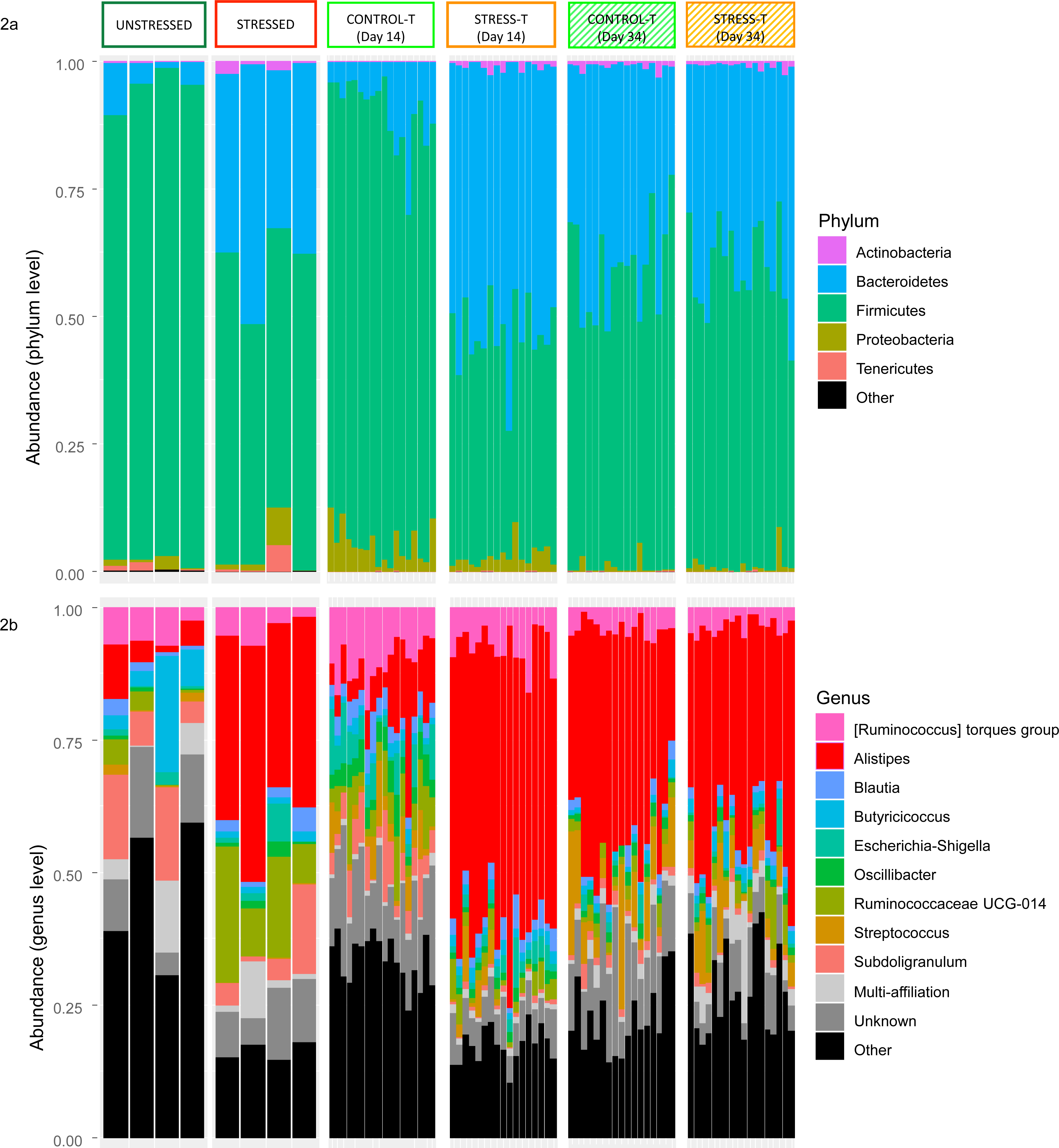
Relative abundance of major bacterial phyla (**a**) and genera (**b**) in the cecal contents of unstressed (n = 4) and stressed (n = 4) quails, CONTROL-T (n = 18) and STRESS-T (n = 17) groups at Day 14 and CONTROL-T (n = 16) and STRESS-T (n = 18) groups at Day 36.

After CMT into germ-free naïve quails, we compared cecal microbiota composition of germ-free recipients colonized with the cecal microbiota of a quail randomly picked either from a group of unstressed quails (CONTROL-T group) or from a group of stressed quails (STRESS- T group).

Cecal contents collected at Day 14 showed differences in the relative abundance of the major phyla between STRESS-T and CONTROL-T groups. Therefore, firstly we assessed overall differences using diversity indexes which revealed a higher cecal microbial diversity for the CONTROL-T quails (Shannon index: 3.94 ± 0.05 *vs* 2.65 ± 0.10, *p* < 0.001 for the CONTROL-T and STRESS-T quails respectively, *p* < 0.001; Inverse Simpson index: 26.08 ± 2.75 vs. 3.96 ± 0.37 for the CONTROL-T and STRESS-T quails respectively, *p* < 0.001; **Figure 1a**). The CONTROL-T and STRESS-T groups compared at OTU level using Bray-Curtis distances revealed high between-group differences and weak within-group differences (*p* < 0.001, **Figure 1b**). Secondly, we observed differential abundances reflecting the cecal microbial composition already observed in the donor quails. At the phylum level, the *Firmicutes* were thus more abundant in the CONTROL-T quails (*p* < 0.001), whereas the *Bacteroidetes* and *Actinobacteria* were more abundant in the STRESS-T quails (*p* < 0.001 and *p* < 0.001 respectively) (**Figure 2a**). At the genus level, differential abundances were found for 40 genera, but only *Alistipes* sp. presented reasonably high relative abundances (> 5% in mean; *p* < 0.001; **Figure 2b**) with higher abundance in the STRESS-T group. This genus was mainly represented by OTU1 and OTU2 (respectively 84.9% and 14.0% of the sequences assigned to *Alistipes* sp.). Thirdly, the functional diversity was inferred from the taxonomic profiling, using the PICRUST2 approach for function (i.e. E.C. numbers) and KEGG (Kyoto Encyclopedia of Genes and Genomes) pathway prediction. The results indicated an average NSTI score of 0.23 ± 0.06 for the 415 OTUs, showing that the predictions were poorly supported for a fraction of the OTUs. However, the NSTI scores were respectively 0.12 and 0.11 for the two OTUs involved in the main differences between the STRESS-T and CONTROL-T groups, OTU1 and OTU2. We found 300 enriched KEGG pathways; among them, 78 pathways presented significant differences between STRESS-T and CONTROL-T groups (**Figure 1c; Supplementary table 1).** In particular, we observed that the microbiota from STRESS-T had a reduction in tryptophan synthesis (EC:4.2.1.20) while its catalysis was enriched compared to CONTROL-T (EC:4.1.99.1; **Figure 1d**), mainly through the *Alistipes* genus which represent 15/26 of the species having tryptophanase.

The cecal contents collected at Day 36 showed few differences between STRESS-T and CONTROL-T groups. The levels of alpha diversities were lower than those observed at Day 14 and were similar between the CONTROL-T and STRESS-T groups (Shannon index: 3.04 ± 0.15 *vs* 3.11 ± 0.12, *p* > 0.10 for the CONTROL-T and STRESS-T quails respectively; Inverse Simpson index: 6.77 ± 1.19 vs. 6.91 ± 1.06 for the CONTROL-T and STRESS-T quails respectively, *p* > 0.10; **Figure 1a**). The Bray-Curtis distances revealed qualitative differences between the groups (*p* < 0.001; **Figure 1b**). However, we did not observe significant alterations in differential abundances at the phylum level (**Figure 2a**) and only one genus presented differential abundances and a reasonably high abundance (5.0 %). This category included all uncertain genera assigned to the *Lachnospiraceae* family (> 5% in mean; *p* < 0.001; **Figure 2b**). In line with this, functional predictions revealed only a few differences between the CONTROL-T and STRESS-T group at Day 36, including two pathways associated with tetrapyrrole biosynthesis (PWY-5189 and PWY-5188; **Figure 1c)** and seven functions mainly associated with these pathways.

### Cecal microbiota transfer from stressed quails to germ-free naïve quails results in increased anxiety-like behavior and impaired spatial and cue-based memory

In a novel environment test, quails from the STRESS-T group spent significantly more time on average in the wall zone, corresponding to where they were introduced and which indicated a fear-induced reduction of exploration (**Figure 3a)**.

**Figure 3:**
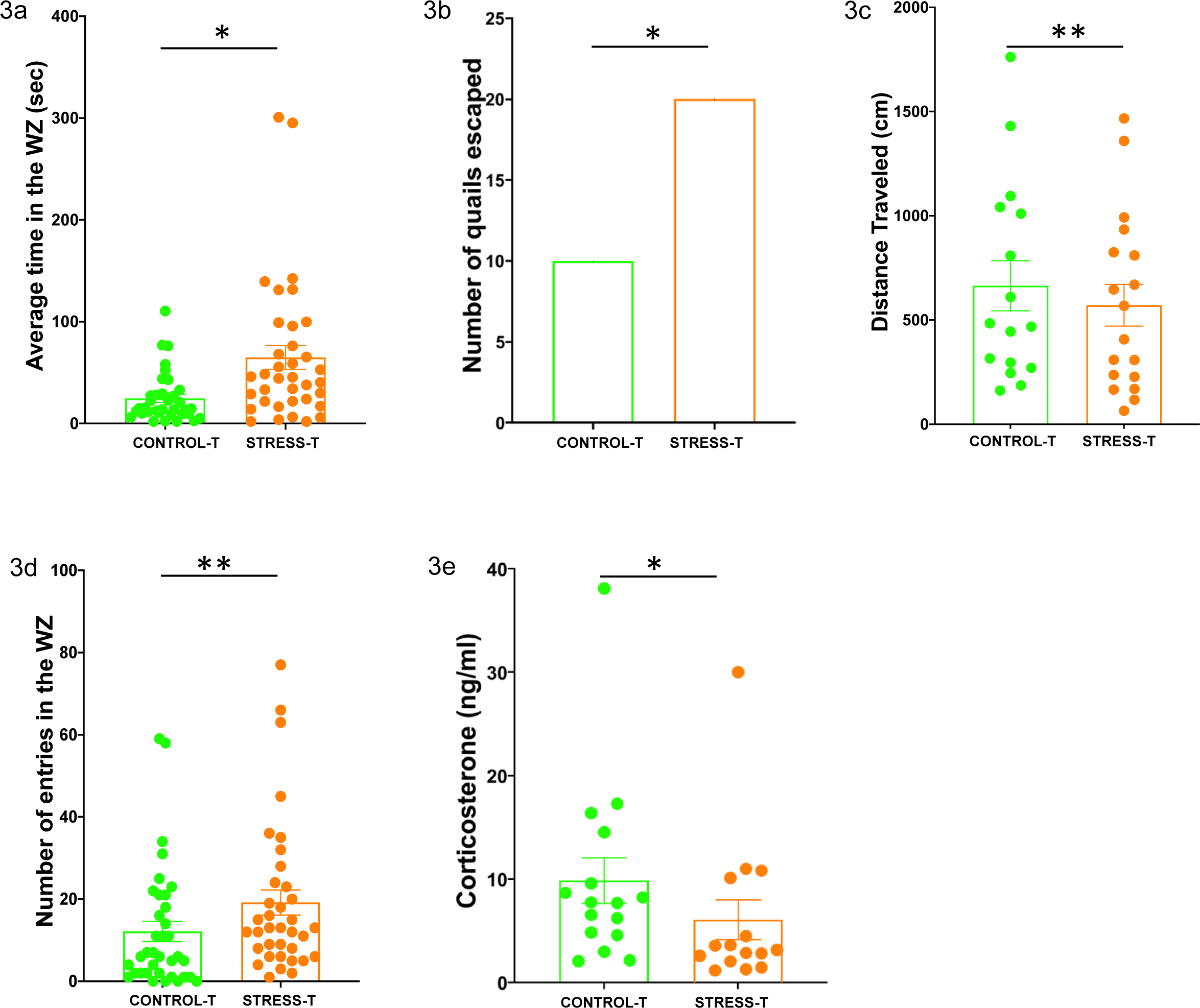
(**a**) Average time spent in the wall zone (WZ) during the novel environment test in the CONTROL-T (n=36) and STRESS-T (n = 36) groups. (**b**) Number of quails that escaped from the device during the novel object test in the CONTROL-T (n = 36) and STRESS-T (n = 36) groups. (**c**) Distance traveled during the open-field test in the CONTROL-T group (n = 16) and the STRESS-T group (n = 18). (**d**) Number of entries in the wall zone (WZ) during the social separation test in the CONTROL-T (n = 36) and STRESS-T (n = 36) groups. (**e**) Basal plasma corticosterone levels on Day 14 in the CONTROL-T group (n = 16) and the STRESS- T group (n = 15). The results are expressed as mean values ± SEM. * *p* < 0.05, ** *p* ≤ 0.01.

Fear of novelty was also investigated using a test that involved introducing a novel object (red plastic ball). The quails that fled far from the object were twice as numerous in the STRESS-T group as in the CONTROL-T group (**Figure 3b**).

In the open-field test, quails of the STRESS-T group traveled significantly shorter distances than those of the CONTROL-T group (**Figure 3c**), which indicates a state of enhanced stress.

When separated from their congeners by a wall in the social separation test, the STRESS group individuals entered significantly more into this wall zone (**Figure 3d)** which reveals increased locomotor activity in this zone, indicating higher anxiety-like behavior in this situation of social isolation.

In the memory testing, quails were individually habituated to the test arena (**Supplementary figure d**) and trained in the spatial learning task before the tests. During the 4 days of training, the quails of both groups learned to find the rewarded cup without treatment effect on the latency to visit the rewarded cup and the number of cups visited before reaching the rewarded cup. Latency to visit the rewarded cup decreased over time independently of the treatment (day effect: **χ^2^** = 24.29, *p <* 0.0001; treatment effect: **χ^2^** = 0.78, *p =* 0.38; interaction day*treatment: **χ^2^** = 0.58, *p* = 0.44, **Figure 4a**) and the number of cups visited before reaching the rewarded cup also decreased without treatment effect (day effect: **χ^2^** = 7.11, *p <* 0.01; treatment effect: **χ^2^** = 0.18, *p =* 0.67; interaction day*treatment: **χ^2^** = 0.36, *p* = 0.54, **Figure 4b**). After training, a test to evaluate spatial memory was performed. During this test, all the cups were unrewarded and quails had to find the previously rewarded cup - at an unchanged location - using only spatial information since all cups had a white cover. This test revealed spatial memory was impaired in the STRESS-T group compared to the CONTROL-T group. Quails of the STRESS-T group tended to take more time (**Figure 4c**) and visited significantly more cups before reaching the location of the previously rewarded cup (**Figure 4d**).

**Figure 4:**
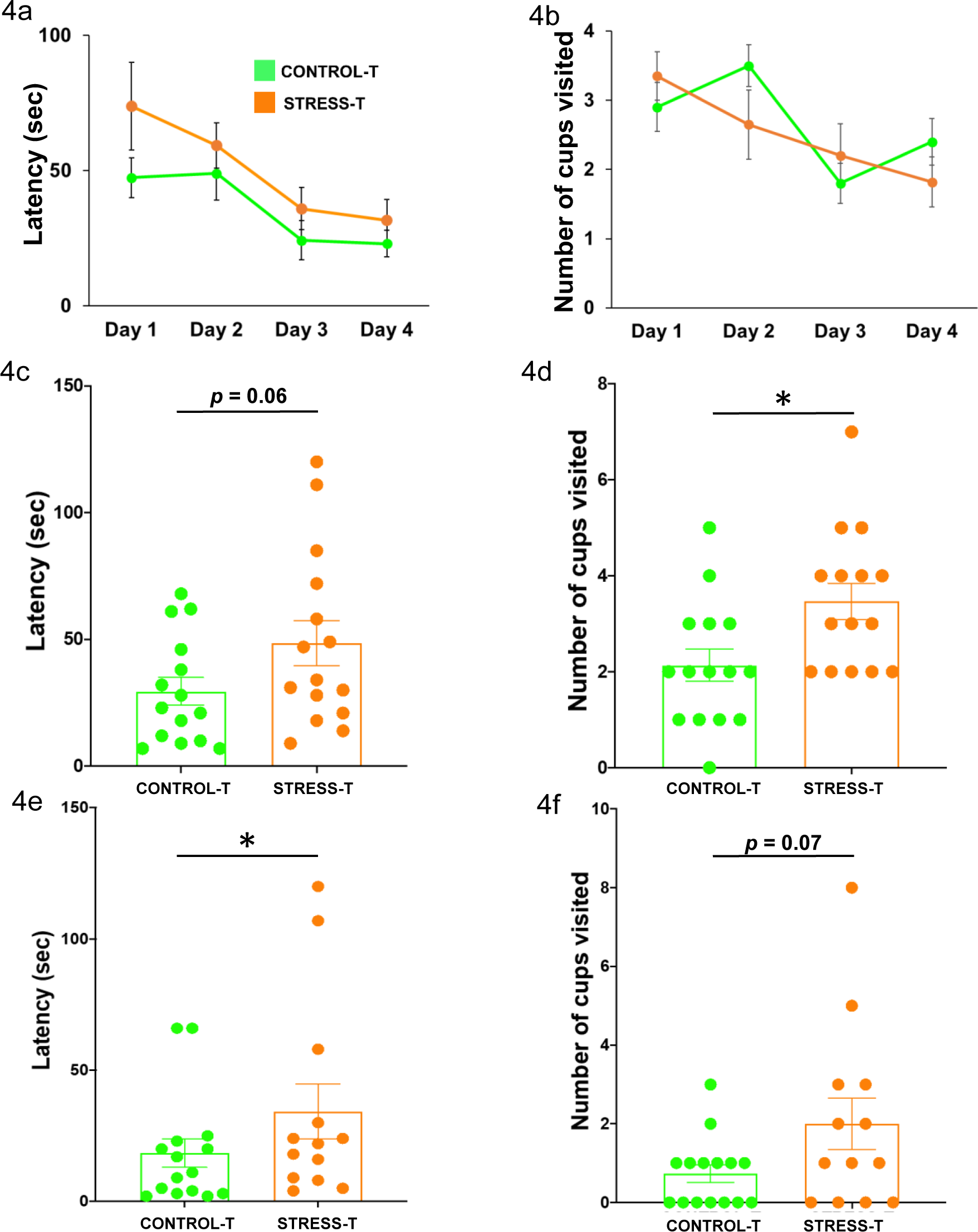
Latency (**a**) and number of cups visited (**b**) before reaching the rewarded cup during the 4 days of training in the CONTROL-T group (n = 15) and the STRESS-T group (n = 15). Latency (**c**) and number of cups visited (**d**) before finding the location of the previous rewarded cup during the spatial test in the CONTROL-T group (n = 15) and the STRESS-T group (n = 15). Latency to reach the cued cup (**e**) and number of cups visited before reaching the cued cup (**f**) in the CONTROL-T group (n = 15) and the STRESS-T group (n = 14) during the cued test. The results are expressed as mean values ± SEM. * *p* < 0.05.

During the cued test, all the cups were unrewarded, the location of the cued cup was modified and we measured whether the individuals looked for the location of the cup usually rewarded (spatial memory) or for the cue that was associated with the reward during training (cue-based memory; black cover). Quails of the STRESS-T group took significantly more time (**Figure 4e**) and tended to make more visits than CONTROL-T quails before reaching the cued cup (**Figure 4f**).

### Microbiota transplantation influences corticosterone levels in the plasma, SCFA concentration in the feces and gene mRNA expression in the brain

We assessed the impact of CMT on plasma corticosterone levels of recipient quails both before and after acute stress. Plasma corticosterone levels at baseline were significantly lower in the STRESS-T group than in the CONTROL-T group (**Figure 3e**). The magnitude of the increase in corticosterone after contention (level induced by stress of contention minus basal level) tended to be higher in STRESS-T quails than in CONTROL-T quails (6.02 ± 1.9 *vs* -1.7 ± 2.8, **χ^2^** = 3.54, *p* = 0.07) and the plasma corticosterone levels obtained after 10 min of restraint stress were not significantly different between the two groups (8.7 ± 0.9 for CONTROL-T group *vs* 11.3 ± 2.5 for STRESS-T group, **χ^2^** = 0.05, *p* > 0.10).

The fermentation activity of the gut microbiota was measured through the quantitative analysis of SCFA contained in fecal contents at Day 6 and Day 20 after CMT. Fecal samples at Day 6 revealed significantly higher concentrations in the CONTROL-T group for caproate, isovalerate and isocaproate (**Supplementary table 2)**. No significant differences in SCFA composition were found at Day 20.

Finally, we assessed CMT effects on gene expression in the hippocampus, the arcopallium and the hypothalamus, which are brain structures involved in cognitive processing, control of fear behavior and regulation of the HPA axis, respectively. In the hippocampus*, CRHR1* expression was significantly lower in quails of the STRESS-T group (**Figure 5a**). No significant differences were found in the arcopallium (**Figure 5b**). The expression of *CRHR1* and *PCNA* were significantly reduced in the hypothalamus of the STRESS-T group compared to the CONTROL-T group (**Figure 5c**).

**Figure 5:**
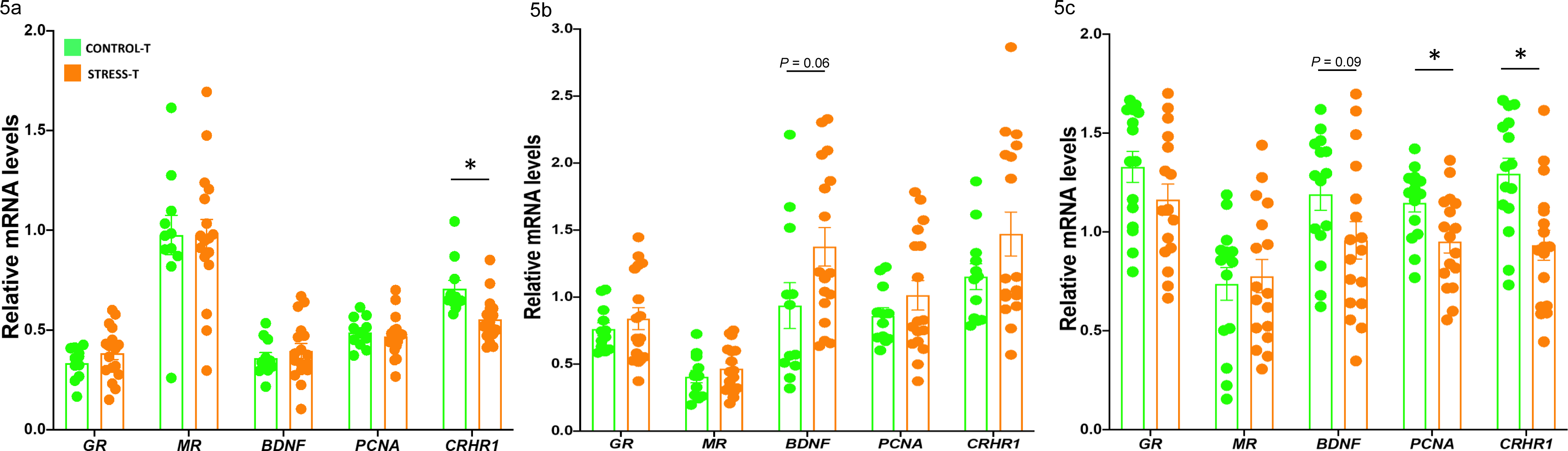
Gene expression in the hippocampus (**a**, CONTROL-T: n = 11, STRESS-T: n = 17), the arcopallium (**b**, CONTROL-T: n = 12, STRESS-T: n = 17) and the hypothalamus (**c**, CONTROL-T: n = 15, STRESS-T: n = 17). The results are expressed as mean values ± SEM. * p < 0.05.

## Discussion

Chronic stress is not only recognized to have a major impact on gut physiology and microbiota composition ^4, 9^ but it is also known to be an important risk factor for brain-related dysfunctions such as memory impairments ^10–15^. Moreover, Chevalier and collaborators ^8^ recently showed that chronic stress and the gut microbiota generate a feedforward loop that contributes to depressive disorders. In the present research, we investigated whether a similar loop system could exist between gut microbiota and memory deficits associated with chronic stress conditions. Using Japanese quails raised in a unique microbial controlled-environment inside isolators, we demonstrated that germ-free host quails receiving cecal microbiota from a donor quail subjected to chronic stress showed clear impairments in memory and also increased anxiety-like behavior when compared to germ-free quails implanted with cecal microbiota from unstressed quail. To induce chronic stress in quails we used a procedure of unpredictable negative stimulations, which has been thoroughly validated in this line of Japanese quail ^39–42^. Although this procedure has profound impacts on behavior and physiology, its potential impact on gut microbiota has not been investigated to date.

The composition of cecal microbiota of quails subjected to the chronic stress procedure differed from that of unstressed quails. The main differences related to one OTU assigned to the *Alistipes* genus (OTU 1; more abundant in stressed quails), which belongs to the *Bacteroidetes* phylum, known to be altered by various forms of stress, including exposure to early-life maternal separation stress ^43^, social stress ^44^ and water-avoidance stress ^45^. *Alistipes* sp. is a genus already known to be favored by various kinds of induced stress in different models. This includes models in which mice were subjected to a water immersion restraint stress ^46^ or mice housed on a grid floor and stressed by this rearing condition ^47^. Interestingly, increased abundance of *Alistipes* sp. has also been found in the gut microbiota of depressive human patients ^48, 49^ and in anorexia nervosa ^50^.

The oral inoculation of the modified microbiota in recipient quails resulted in a higher anxiety level than in quails inoculated with the microbiota from unstressed quail. The rigorous use of germ-free chicks and controlled conditions in isolators demonstrates that individuals of the STRESS-T group were more anxious as they displayed an overall decrease in exploration in the novel environment test and the open-field test and increased activity during social isolation. This increase in anxiety-like behavior mimics the one reported in quails subjected to the chronic stress procedure ^40, 51^. The higher anxiety-like behavior of STRESS-T quails in response to novelty was confirmed by the novel object test during which the quails escaped twice as much in the STRESS-T group compared to the CONTROL-T group. Again, this result is in line with the increased neophobia described in chronic stress quails ^41, 52^ and further strengthens the implication of gut microbiota in neophobia responses that we recently characterized in this quail line ^34^.

Our study showed that colonization with the cecal microbiota from a stressed individual affects spatial memory and cue-based memory. During the training session, the quails of both groups learned the task similarly, which allows us to interpret the test responses in terms of specific memory capacity with no bias of motivation, vision ability or learning.

Spatial memory, which consists in relating positions of visual cues in the environment, is indeed a privileged target of chronic stress in mammals and also in birds ^53, 54^. Unlike spatial memory, which is a form of explicit memory, cue-based memory is an implicit memory system based on a simple cue-response association ^55, 56^. Previous studies have shown correlations between modifications of the gut microbiota and impaired memory performances ^16^. However, no study has established a causal link between the alterations in gut microbiota induced by chronic stress and memory deficits. The CMT protocol we used enables us to demonstrate for the first time the causal role of the gut microbiota in stress-induced memory impairments by showing that an altered gut microbiota alone is able to induce the negative effects of chronic stress on spatial and cue-based memory (**Figure 6**). This pivotal result is in line with a recent study that showed that a transfer of gut microbiota from old mice to young mice was sufficient to reproduce the cognitive decline associated with aging ^57^ and suggests that memory impairments are mediated by gut microbiota in many cases. The results of the CMT protocol provide evidence of a feed-forward loop system linking the microbiota-gut-brain axis to stress and memory function (**Figure 6**). This evidence suggests that future research should target gut microbiota composition and not only neurobiological pathways to prevent stress-induced memory alterations.

**Figure 6:**
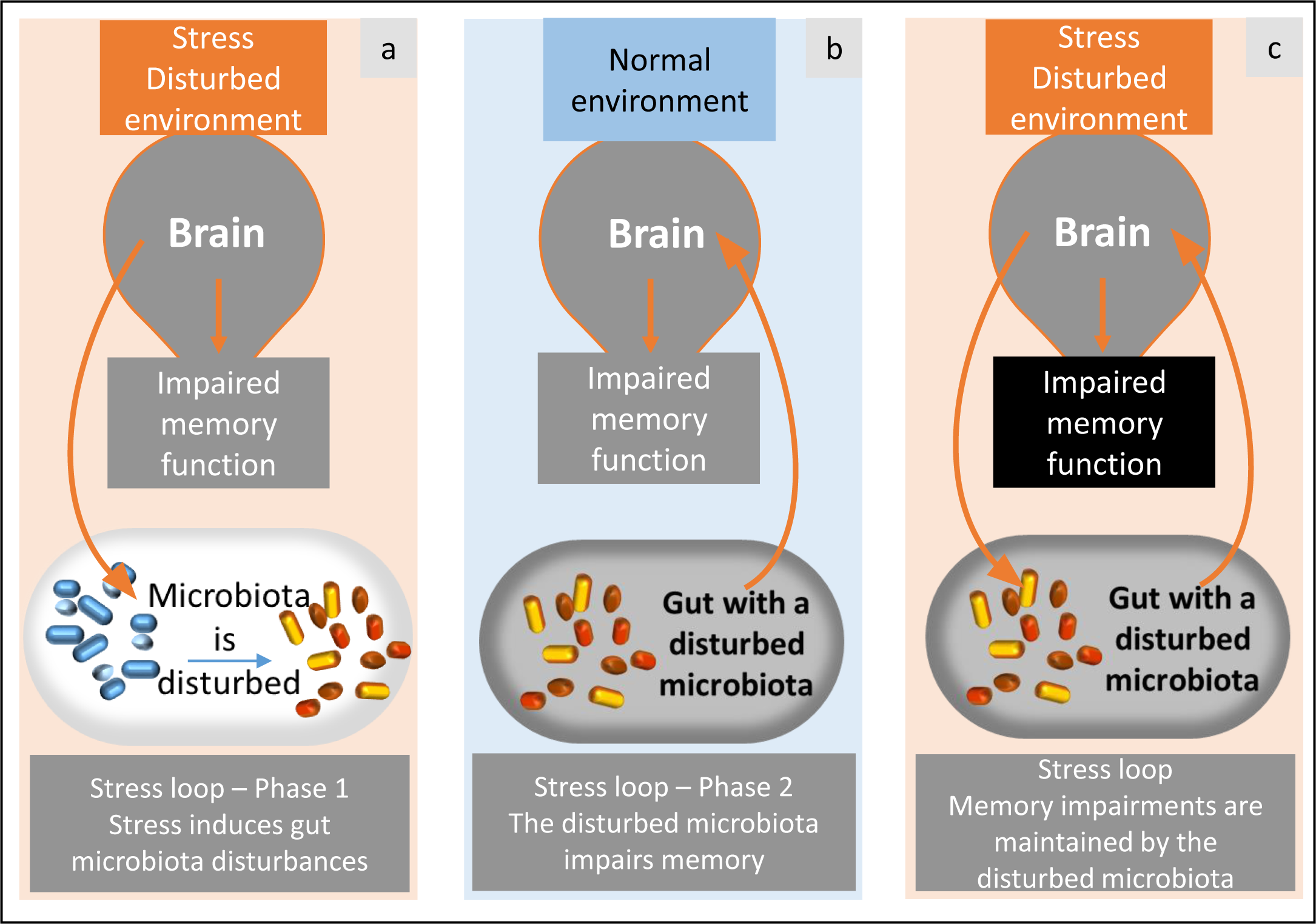
A vicious loop *via* gut microbiota. Stress and disturbed environment conditions (**a**) induce alterations in the gut microbiota. This disturbed gut microbiota is able to induce impaired memory by itself, even if the environment is not stressful (demonstration in the present study, **b**). As a consequence, the gut microbiota and brain would result in a loop that maintains memory impairments (**c**).

Furthermore, our data revealed differential regulation of several genes in the brain according to the cecal microbiota used for colonization. In the hippocampus and the hypothalamus, colonization with the cecal microbiota from a stressed individual reduced *CRHR1* expression and the level of plasma corticosterone. *CRHR1* is an essential regulator of the HPA axis; its hippocampal expression has also been shown to be reduced by maternal separation in mice ^58^ and *CRHR1*-deficient mice are unable to mount a corticosterone response to stress ^59^. Moreover, this reduction in plasma corticosterone levels under basal conditions has previously been described in European starling ^60, 61^ and in this line of quail subjected to a chronic stress procedure ^40^, whereas more acute stress increases corticosterone levels in this species ^38^. We found reduced *PCNA* expression in the hypothalamus of the STRESS-T group, which suggests decreased cell proliferation. We also noted that in the STRESS-T group *BDNF* expression tended to reduce or increase in the hypothalamus and arcopallium respectively, suggesting changes in brain plasticity mechanisms. However, we did not detect any significant differences in mRNA expression levels in any of the three brain structures for the nuclear receptors *GR* and *MR* involved in the negative feedback of the HPA axis.

We looked at the cecal microbiota composition to understand more clearly whether microbiota transfer can modulate anxiety-like behavior and memory performance. As expected, the cecal microbiota transfer also led to different cecal microbiota composition in the recipient quails which can explain the behavioral, cognitive and physiological differences observed. At Day 14, cecal contents of STRESS-T quails showed lower microbial alpha diversity, a lower abundance of *Firmicutes* and a higher abundance of *Actinobacteria* and *Bacteroidetes* than cecal contents of CONTROL-T quails. Interestingly, these results are very similar to those observed in stressed or unstressed donors, which suggests successful microbiota transfer. In addition, the greatest differences in OTUs between the two groups were assigned to the *Alistipes* genus with higher abundance in the STRESS-T than CONTROL group. As previously mentioned, the *Alistipes* genus has already been linked to stress and depression in mice and humans ^46–48^ and could perhaps serve as a biomarker of stress. An increasing body of literature supports different explanations for the mechanisms by which *Alistipes* could play a role in the MGBA ^62^. Detrimental effects of *Alistipes* would be related to the permeability of the gut induced by microbial dysbiosis which allows molecules such as lipopolysaccharides (LPS) to enter into the bloodstream leading to neuroinflammation and behavioral alterations. The resulting inflammatory cytokine production in the central nervous system impairs the synthesis of neuropeptides associated with brain-related disorders, including depression ^63^. The explanation that best fits our results and our microbiome functional analysis is that of the tryptophan amino acid (Trp) pathway and the serotonergic system since *Alistipes* species are indole-positive and possess the tryptophanase enzyme which directly produces indole from Trp and may lead to a disruption of the serotonergic balance in the host ^62, 64, 65^. Since the link between the serotonergic system and HPA axis is well-recognized now ^66^, the alterations of behavior and HPA axis activity found in our study could be explained by the high abundance of *Alistipes* species and their action on the Trp pathway. In addition, our results on the tryptophanase function in the gut microbiota are in line with several studies showing a link between Trp metabolism in the gut and behavioral changes during chronic stress ^67–69^. These findings in *Alistipes* make this bacterial genus an important candidate in the interaction between the gut microbiota and the stress system and corroborate the implication of microbial Trp metabolism in behavioral and neurological stress-induced changes.

The fermentation activity of the gut microbiota assessed through the quantitative analysis of the SCFA contained in fecal contents showed higher concentrations of caproate, isocaproate and isovalerate, but lower proportions of acetate in CONTROL-T quails. Together, these data showed that CONTROL-T and STRESS-T quails had different gut microbiota fermentation activities, which strongly supports the involvement of SCFA in the microbiota-gut-brain communication in vertebrates ^70–72^ and is in line with recent results showing that microbiota changes induced by chronic stress affect lipid metabolism and the generation of endocannabinoids ^8^.

At the end of the experiment after 36 days, there were fewer differences in cecal microbiota between STRESS-T and CONTROL-T groups. This could reflect an age effect and an evolution of the cecal microbiota with time or a change of environment during the memory test procedure. However, both groups still showed differences in terms of anxiety-like behavior, cognition and gene expression in brain structures. This may imply that there is a critical period in early life during which the initially implanted microbiota would have irreversible consequences on cerebral, behavioral and cognitive development even after re-colonization with a different microbiota. This hypothesis is supported by several rodent studies and would imply that a critical period may exist in all vertebrates ^73, 74^. These long-term effects of the microbiota suggest that the origin of certain cognitive disorders should not only be investigated in the gut microbiota present at the time of the onset of the disorders, but also in previous intestinal changes. Interestingly, studies aimed at investigating a potential correlation between autism spectrum disorders and antibiotic treatments received at a young age revealed an imbalance in the composition of the gut microbiota ^75^. This suggests that the prevention of memory alterations due to stress must target an immediate return to a state of equilibrium in the composition of the microbiota in order to avoid possible long-term effects.

## Conclusions

In conclusion, we showed that gut microbiota alone is sufficient to mimic stress effects on cognition and impair memory abilities. These data substantiate the existence of a stress loop connecting the gut to memory development that implicates the gut microbiota as a component that has to be considered in greater depth in future studies on stress processes. Interestingly, our findings add more evidence to the role of *Alistipes* genus as a potential biomarker of stress in vertebrates because of its link with the tryptophan metabolism pathway. These data suggest that maintaining a healthy microbiota could help alleviate memory impairments linked to chronic stress.

## Materials and Methods

All the animal care procedures were carried out in accordance with the guidelines set by the European Community’s Council Directive (DIRECTIVE 2010/63/UE) and with French legislation on animal research. The protocol was approved by the French Ministry of education, higher education and research (Ministère de l’Education Nationale, de l’Enseignement Supérieur et de la Recherche) under the protocol N° APAFIS# 201707131037724.V3 - 10607. The principles of reduction, replacement and refinement were implemented in all the experiments. We used a line genetically selected for its long tonic immobility duration and therefore a high level of emotional reactivity (E+) ^76^. Emotional reactivity is characterized by behavioral and physiological responses to a challenging situation ^77^.

The timing plan of the experiment is summarized in the **Supplementary figure a**.

### Chronic mild stress procedure in donors and microbiota sample collection

The quail chicks used to provide the stressed and unstressed donors were hatched on the farm of the Experimental Poultry Facility (UE PEAT, INRAE, 2018. DOI: 10.15454/1.5572326250887292E12). At 15th day of age, male quails were divided in two groups, i.e. unstressed and stressed groups. From the age of 17 to 40 days, quails from the stressed group were exposed to unpredictable repeated negative stimuli (confinement in a corner of the home cage, disturbances in the home cage, cage shaking, noises, crowding, novel environment, transport) four times per day and once per night while quails from the unstressed group were just visited by a human four times per day according to the procedure described by Favreau-Peigné et al.^32, 33^. Each negative stimulus lasted 30 min, continuously or not. Negative stimuli and visits occurred at random times and a given stimulus was never used twice per day in order to increase unpredictability and decrease animal habituation to the stress procedure.

Cecal contents were collected from 4 adult males of the E+ line which had previously been subjected to the chronic stress procedure described ^32, 33^ during both night and day for 21 days (stressed quails) and from 4 adult males of the same line which had not been stressed (unstressed quails). Both groups of quails aged 6 weeks old were housed under the same conditions and fed the same diet. For cecal content collection, both ceca were opened. Their contents were gently removed to obtain only the contents and not the mucosa and then mixed in 500 μL of sterile glycerol + cysteine. All the cecal contents were collected in a clean room with autoclaved tools under a sterile biological safety cabinet and were then stored at – 80°C.

### Production of germ-free quail recipients

The eggs were collected every 90 min from females from the same line reared under conventional conditions and that were not exposed to any stress procedure. The eggs were disinfected following the procedure previously described ^34^ in the facilities of the PFIE, INRA, 2018 (Infectiology of Farm, Model and Wild Animals Facility, https://doi.org/10.15454/1.5572352821559333E12). Briefly, eggs were disinfected with potassium permanganate (10 g) and a solution of formalin + milliQ water for 90 min and in Divosan-plus (Diversey France SAS) 1.5% for 1 min before incubation for 14 days. After a third disinfection with a spray of Divosan 1.5% for 30 s, eggs were rinsed with autoclaved water for 16.5 min.

### Animals and housing

Disinfected eggs were placed in sterile isolators in the facilities of the PFIE and germ-free chicks hatched in the following days. Control for germ-free status was performed through anaerobic and aerobic culture of freshly voided fecal samples (in resazurin thioglycolate, serum enriched BHI and blood agar and incubated at 25°C and 35°C). Two days after hatching (Day 2), the chicks were transferred to the breeding room which housed six large sterile isolators of identical dimensions and organization (**Supplementary figure b;** see Kraimi et al. ^34^). Using the Polymerase Chain Reaction (PCR) method on down feathers for sexing Japanese quail at hatching ^78^, 6 females and 6 males were kept in each isolator for the experiment. All quails were given free access to γ-irradiated (45kGy, Scientific Animal Food and Engineering, Augy, France) feed pellets (metabolizable energy: 12.2 MJ; Crude Protein: 204 g/kg) and autoclaved water. The temperature in the isolators was set at 40-38°C the first days and was gradually reduced to 28°C at 14 days of age. The light phase was 24 hours until Day 4 and this was progressively reduced by 1 hour per day until reaching a minimum of 10 hours of light per day after 18 days. The living environment of the chicks in the isolators was enriched by wood shaving dust baths and by placing previously sterilized new objects (plastic or glass balls) on successive days. In order to avoid mating in the isolator and excessive stocking density, female quails were eliminated at Day 14 and only males were conserved for plasma corticosterone assays and memory tests.

### Bacterial inoculation

On the day of transfer to the isolators (Day 2), we colonized the chicks of three of the isolators (group STRESS-T) with the cecal microbiota from a unique donor male randomly chosen from the stressed quails. This was done to maintain a balanced ecosystem. The chicks from the three other isolators (group CONTROL-T) were colonized with the cecal microbiota from another donor male randomly chosen from the unstressed quails. For each group, the cecal content was thawed and diluted aerobically in 12 mL of sterile physiological saline. Each chick was colonized by oral gavage with 100 µL of this microbiota suspension.

The cecal contents of recipient females at Day 14 and recipient males at Day 36 were also collected for analysis of microbiota composition following the same procedure as described above.

### Microbiota composition analysis

#### Microbial DNA extraction and 16S rRNA gene amplification

Microbial DNA extraction was performed using the QIAamp DNA mini-kit (ref #51306, Qiagen Inc., Courtaboeuf, France) following the procedure previously described^36^. Briefly, 25 mg of thawed digestive content were mixed in 1 mL of lysis buffer and homogenized at maximum speed (frequency 30.sec-1) with 0.4 g of sterile zirconium beads in a tissuelyser mixer Retsch MM400 for 3 min, followed by heating at 70°C for 15 min. After a centrifugation at 16 000 g and 4°C for 5 min, the supernatant was conserved at ambient temperature and 300 μL lysis buffer were added. The homogenization steps were repeated on the pellet and followed by a second centrifugation (5 min, 16 000 g, 4°C). The two supernatants were pooled and homogenized for the DNA purification and filtration step. Proteinase K and AL buffer were added to the supernatants and the mix was heated at 70°C for 10 min to remove proteins. The sample was transferred to a tube containing pure ethanol for a purification step using a QIAamp column as described by the manufacturer. The sample was then eluted in buffer AE (10 mM Tris-Cl; 0.5 mM EDTA; pH 9.0; Qiagen Inc., Courtaboeuf, France). DNA purity was assessed with a NanoDrop spectrophotometer based on the 260/280 and 260/230 O.D. ratios.

PCR amplification of the bacterial 16S rRNA gene on DNA extracts was carried out using the primers designed to amplify from highly V4-V5 conserved regions (Forward: CTTTCCCTACACGACGCTCTTCCGATCTGTGYCAGCMGCCGCGGTA; Reverse: GGAGTTCAGACGTGTGCTCTTCCGATCTCCCCGYCAATTCMTTTRAGT). The PCR program used consisted of an initial denaturation at 94°C for 2 min followed by 30 cycles of 94°C for 60 s, 65°C for 40 s, 72°C for 30 s and a final extension step of 72°C for 10 min. PCR product size was checked with 2% agarose gel electrophoresis before the sequencing step.

#### 16S rRNA gene sequencing

V4-V5 region full length reads were obtained using Illumina Miseq 250-bp paired end reads. The resulting PCR products were purified and loaded onto the Illumina MiSeq cartridge in accordance with the manufacturer’s instructions. Using PhiX control following the manufacturer instructions, the quality of the run was checked internally and with the help of the previously integrated index, each pair-end sequence was assigned to its sample. Each pair-end sequence was assembled using Flash software with at least a 10-bp overlap between the forward and reverse sequences, which allows 10% mismatch. The absence of contamination was checked with a negative control during the PCR (water as template). The corresponding sequences were uploaded on NCBI with the access number (PRJNA527873).

#### Anxiety-like behavior tests

Each isolator was divided into two equal areas using an opaque separating wall. One area was dedicated to rearing, containing the feed and water, while the other half was specifically used for the behavioral tests (see Kraimi et al. ^34^). All behavioral tests were recorded with a camera fixed above each isolator.

#### Novel environment test

On Day 7, in order to assess the anxiety-like behavior in a novel environment we introduced the quails in groups of three (to limit the social isolation component) in the test area for the first time for 5 min and The Observer XT (version 12.5) software was used to measure for each individual the average time spent (time/number of entries) in the wall zone of the test area separated from the rearing area by an opaque wall (**Supplementary figure c**). The quails were initially placed in the wall zone and the time spent in the other parts of the test area was considered as exploration activity.

#### Novel object test

On Day 13, we measured the behavioral reactions of quails in the presence of a novel object. Each quail was placed for 5 min in the test area in a sterilized white plastic corridor which contained a sterilized red plastic ball. The number of quails attempting to escape from the object was counted as an indication of fear.

#### Open-field test

After the memory tests, male quails were subjected to an open-field test outside the isolators to assess anxiety-like behavior to a novel environment. The test was carried out in a clean sterilized room. Quails were removed individually from the isolator and carried in a transport box where they were left for 5 min before the test in order to calm down and limit fear reactions linked to the removal from the isolator. The open-field device was a square arena (80 cm × 80 cm × 29 cm) made of wood with a floor with a yellow waterproof plastic surface under 50 lux light conditions and a camera fixed above the area. Each quail was placed in the center of the open-field and allowed to freely explore the test arena for 5 min. Using the Ethovision XT tracking software (version 7.1), we recorded the locomotor activity (total distance traveled). At the end of each test session, the quail was returned to its isolator using a transport box and the test arena was disinfected.

#### Social separation test

On Day 12 we measured the anxiety-like behavior of the quails during a period of separation from their congeners. This test is an adaptation of the well-known social isolation test because under our conditions, the quails could hear their congeners on the other side of the wall. The quails were individually placed in the test area for 5 min. The Ethovision XT (version 7.1) software was used to record the number of times the quail entered in the wall zone of the test area previously described (**Supplementary figure c**). This parameter represents a good indicator of the anxiety level of the quails because the more agitated a quail is in an attempt to reach its congeners reflects an increasing degree of anxiety ^79^.

#### Plasma corticosterone levels

Plasma corticosterone levels of male quails were measured on Day 14 for the basal value and after a stress (restraint in a crush cage for 10 min) on Day 15. For this purpose, each quail was gently removed from the isolator and transported to a quiet place with no other birds around using a transport box disinfected between each quail. Both basal and acute stress blood samples were collected by jugular puncture into a tube containing EDTA. Sampling alternated between CONTROL-T and STRESS-T group and was carried out during the first 6 h of the light phase (from 9 to 13 am) for both days. Following centrifugation at 4000 g for 10 min at 4°C, plasma samples were separated and stored at −80°C until measurement. Corticosterone was measured using a chemiluminescent immunoassay kit (Corticosterone chemiluminescent immunoassay kit, Arbor Assays, Michigan, USA). Plasma was diluted 1:100 and the measurements were performed with the mean of two replicates. The intra-assay coefficients of variation were 5.9% and 11.0% at 207.7 pg/mL and 64.37 pg/mL, respectively. The inter-assay coefficients of variation were 11.3% and 15.1% at 199.6 pg/mL and 55.6 pg/mL, respectively. The assay sensitivity was 6.71 pg/mL. Three quails of the STRESS-T group and one quail of the CONTROL-T group were removed from the analysis because of problems during sampling.

#### Memory tests

Memory tests were performed on males with 18 quails in the STRESS-T group and 16 quails in the CONTROL-T group (two quails were eliminated from this group due to sexing error). Because of the large size of the memory test device, the entire memory part was conducted outside the isolator in clean disinfected rooms and the quails were always manipulated by the same experimenter with gloves. The behavioral parameters were scored manually directly with a camera (Sony DCR-SR58E) placed above the arena and linked to a computer.

#### Familiarization to mealworms

From Day 17 to Day 19, a familiarization process began with cups and mealworms. Six times per day, each quail was individually removed from the isolator and placed in a transport box with an opaque ceramic cup (6 x 7 cm) containing three live mealworms. The familiarization phase ended when the quails had eaten at least one mealworm in the cup, and the habituation phase began.

#### Habituation

Thirty min before each habituation session, quails were removed from the isolators in groups of three and placed in a pre-test box (one box per isolator, 50 x 40 x 30 cm, with wood shavings) without access to food in order to enhance food motivation. During the 3 days of habituation, quails were individually introduced once a day into the center of an octagonal arena surrounded by walls (50 cm high). The floor was covered by yellow waterproof linoleum under 20 lux light conditions in the center. The arena was surrounded by a blue curtain preventing escape attempts. Four black visual cues were placed on the curtain and 4 others on the walls of the arena. Eight opaque ceramic cups similar to those used during familiarization were placed in the arena. Four cups were covered with black paper and the other four with white paper. Each cup contained a live mealworm and the position of white and black cups were randomly moved each day of habituation and for each quail. Quails were allowed to freely explore the arena and the cups until they found and ate all the mealworms or after a maximum test duration of 600 s. The arena was disinfected after each animal session. The number of mealworms eaten from white and black cups was scored after each habituation session. The habituation phase ended when quails had eaten 6 to 8 mealworms on average per session without a significant difference in performance between the two treatments and after this the training phase started. Three quails of the STRESS-T group which did not approach the cups during the 3 days of habituation were removed from the analysis.

#### Training

As previously, quails were removed from the isolators 30 min before each training session, in groups of three and placed in the pre-test box without access to food in order to enhance food motivation. In the training phase, seven cups were covered with white paper and only the rewarded cup was covered with black paper (**Supplementary figure d**). The rewarded cup contained two to three live mealworms and the location of this cup remained the same throughout the whole training period. Quails underwent two training trials per day in this test design with an interval of 30 min between each trial. Finding the reward is a task which can be solved by quails by either learning that the black cup contains the reward (cue-based memory) or by learning the spatial location of the rewarded cup (spatial memory). Quails were placed in the arena at one of three different randomly distributed entry points. The trial was stopped when quails found the mealworms in the rewarded cup or after a maximum test duration of 300 s. As in the habituation phase, the arena was disinfected between each animal. Between each trial, the quail was returned to its box with its congeners and wood shavings. The latency and the number of cups visited before finding the rewarded cup were recorded for each trial. After 4 consecutive days we stopped the training phase when all the quails took on average less than 35 s and made fewer than 3 mistakes before reaching the black rewarded cup without a significant difference of performance between the two treatments.

#### Probe tests

The day following the last training trial, quails completed two different tests (spatial and cued test). In both of these, no mealworms were placed in the cups to avoid any olfactory cue. In the first, the spatial test, all the cups were white to assess whether quails used their spatial memory to locate the position of the rewarded cup (spatial cup). The tested quail was introduced at a different entry than the three used for the training phase and was allowed to explore the arena freely for 2 min. The latency and the number of cup visits before finding the spatial cup (used as indicator of memory errors) were recorded. In order to prevent a potential influence of one test on the other, the second test was performed 3 days after the spatial test (2 consecutive days of training between the two tests with the rewarded cup in the same location as in the first training period). This second cued test was a displacement test in which the black cup was placed in a different position from that of the training period. This test was used to determine the memory system engaged to solve the task: if a quail went first to the spatial cup from the previous test, it indicates that it used a dominant spatial strategy and if it visited the black cup (the cued cup), it indicates that a dominant cue-based strategy based on the cup color was used ^80^. A new entry point equidistant from the spatial cup and the cued cup was used. The latency and the number of cup visits before finding the spatial or the cued cup were also recorded in this test.

### Gene expression in the brain

#### Tissue collection

At Day 36, quails were decapitated post-euthanasia (administration of 0.3 mL of Vetoquinol Dolethal 182.2 mg/mL in the occipital sinus) and brains quickly removed for dissection of the hippocampus, arcopallium and hypothalamus regions according to the quail brain atlas ^81^. All the samples were immediately deep-frozen in liquid nitrogen and then stored at -80°C until analysis.

#### RNA extraction, reverse transcription and real-time polymerase chain reaction (qPCR)

Total RNA was extracted from frozen brain tissue (hippocampus, arcopallium, hypothalamus) using Trizol Reagent (Sigma) following the manufacturer’s instructions. Briefly, 1 mL of Trizol Reagent was added to each sample and homogenized, 200 µL of chloroform (AnalaR NORMAPUR) were added, and the aqueous phase was precipitated with 500 µL of isopropanol (Carlo Erba Reagents). RNA pellets were rinsed with ethanol 70% (Carlo Erba Reagents) and dissolved in RNase-free water. Concentration and purity of individual RNA samples were assessed with NanoDrop 2000 (ThermoScientific) (260/280 O.D. ratios) and integrity was checked using agarose gel (2%) electrophoresis. All RNA samples were stored at -80°C.

For each sample, 1µg of total RNA was reverse transcribed to cDNA using Omniscript Reverse Transcription Kit (Qiagen) and OligodT primers (10 µM; Eurofins) in a final volume of 20 µL following the manufacturer’s recommendations, then treated with RNase inhibitor (Promega).

Primers were designed using Japanese quail sequences (*Coturnix japonica* 2.0), in exonic regions common to all predicted variants, with Primer-BLAST NCBI (https://www.ncbi.nlm.nih.gov/tools/primer-blast/) and synthesized by Eurogentec. Primers sequences are specified in **Supplementary table 3**. The absence of primer secondary structure was analyzed using OligoEvaluator (Sigma-Aldrich, http://www.oligoevaluator.com/).

Quantitative PCR was performed on CFX-96 Real-Time PCR Detection System (Bio-Rad) using SsoAdvanced Universal SYBR Green Supermix (Bio-Rad) following the cycling conditions consisting of 95°C for 5 min, 39 cycles: 10 s at 95°C / 15 s at 60°C / 10 s at 72°C, and 95°C for 10 min. A melting curve stage was added to ensure the presence of a unique amplicon. All the qPCR reactions were run in triplicate. The optimal cDNA dilution and calibration curves for each primer (10µM) were established using a serial dilution (from 1/4 to 1/64) of a mix of individual cDNA from each group. Efficiencies accepted were ranging from 95 to 105%. Stability of housekeeping gene ^82^ expression between groups was not the same for each brain region. Consequently, hippocampus mRNA levels were normalized to *SUZ12* expression, arcopallium mRNA levels to *GAPDH* and *PGK1* expression, hypothalamus mRNA levels to *GAPDH*, *PGK1* and *β-actin* expression. Normalization was achieved with the 2^−*ΔΔCt*^ method and the normalization factor calculated from geNorm software (version 3.5, https://genorm.cmgg.be/).

#### Short-chain fatty acid (SCFA) analysis in fecal samples

Fecal contents from male quails were collected individually inside the isolators at Day 6 and 20 by placing the quail in a box with a γ-irradiated plastic sheet on its bottom (45kGy, Scientific Animal Food and Engineering, Augy, France). The plastic sheet was changed between each quail to obtain individual samples. All the samples were stored at – 80°C before SCFA analysis. Samples were water extracted and proteins were precipitated with phosphotungstic acid. A volume of 0.1 µL supernatant fraction was analyzed for SCFA on a gas–liquid chromatograph (Autosystem XL; Perkin Elmer, Saint-Quentin-en-Yvelines, France) equipped with a split-splitless injector, a flame-ionization detector and a capillary column (15 m x 0.53 mm, 0.5µm) impregnated with SP 1000 (FSCAP Nukol; Supelco, Saint-Quentin-Fallavier, France). Carrier gas (Hydrogen) flow rate was 10 mL/min and inlet, column and detector temperatures were 200°C, 100°C and 240°C, respectively. 2-Ethylbutyrate was used as the internal standard ^83^. Samples were analyzed in duplicate. Data were collected and peaks integrated using the Turbochrom v. 6 software (Perkin Elmer, Courtaboeuf, France).

### Statistical analysis

The results are presented as means ± SEM. The significance level was set at p ≤ 0.05 and 0.05 < *p* ≤ 0.10 was considered as a trend. All statistical analyses were performed with R (version 3.5.1) and RStudio software (version 1.1.463).

The 16S rRNA gene sequences were clustered in operational taxonomic units (OTU) using Swarm. We performed additional filtering steps based on abundance criteria (i.e. the OTUs should be found in at least 3 samples; the minimum abundance threshold was set to 100 sequences across all samples), chimera detection, and comparison to a PhiX contaminant database: this was performed using the dedicated tools implemented in FROGS. Lastly, the OTU taxonomic assignation was performed by a BLAST comparison against the silva 16S 132 database followed by a careful manual curation. We only kept the OTUs presenting an abundance above 0.001 (for a total of 110 OTUs).

Inverse Simpson and Shannon α-diversity indexes were calculated using the R-package vegan v. 2.5-6. Bray-Curtis distances were calculated and tested with ADONIS for significance ^84^ and summarized by multidimensional. Differential abundances at the genus level among E+ stressed quails, E+ unstressed quails, STRESS-T and CONTROL-T groups were assessed using Welsch t-tests corrected using Bonferonni’s approach. The computations were performed using STAMP v 2.1.3.

Finally, the samples were compared using PICRUST2 v. 2.2.0_b in order to infer microbial gene content from 16S rRNA gene data and associated enrichment of metabolic pathways ^85^. The distribution of pathway abundances was visualized by PCA, using STAMP v 2.1.3; differential pathway abundances were assessed using Welch’s t-tests corrected using Bonferonni’s method.

Behavioral data were analyzed using generalized linear mixed models (GLMM; package ‘lme4’ v 1.1-19) with group (STRESS-T or CONTROL-T), sex and interaction between group and sex as the fixed effects and the order in which the quails were tested as the random effect. In the novel environment test, where three quails were tested together, the trio number was used as the random effect. In the case of repeated measures as in the habituation and learning phase of the memory tests, group and day were used as the fixed effects with the order of passage as the random effect. GLMMs were used when data was not normally distributed: a GLMM with Gamma errors was used for the total distance traveled, the latencies and the time spent in the various zones during the different behavioral tests. A GLMM with Poisson errors was used to compare the number of entries in the different zones of the tests and the number of cups visited in the memory tests. During the novel object test, we compared the number of quails that escaped in each group using a Chi2 test.

Corticosterone data were first log-transformed and then tested using a generalized linear model (GLMM; package ‘lme4’ v 1.1-19) with group as the main factor and the order of collection as the random factor.

Gene expression data and SCFA concentration were also analyzed with generalized linear mixed models (GLMM; package ‘lme4’ v 1.1-19) with Gamma law and the group as the fixed effect.

## Declarations

### Availability of data and materials

The datasets used during the current study are available from the corresponding author on reasonable request and they are available at https://doi.org/10.15454/JYITK4. Illumina sequence data have been deposited at National Center for Biotechnology Information (NCBI), under the BioProject PRJNA 527873.

### Author’s contributions

CL and NK designed the study with the help of LC and SR. CL and NK performed the experiments with the technical help of JL, CP, PC, TC and P. Cousin. FL performed the chronic stress procedure. The DNA microbial extraction and PCR steps were carried out by KG and CD. OZ was in charge of 16S rRNA gene sequencing and FK performed the statistical analysis of microbiota data. AF and MPM were in charge of plasma corticosterone measures. JL, NK, AVC, VC and HD performed RNA extraction and qPCR on brains. CP performed SCFA analysis. NK and CL wrote the manuscript. All the authors reviewed and approved the final manuscript.

## Acknowledgements

We thank Sandrine Rivière and Michael Troquet for providing germ-free eggs and conducting blood sampling (UE PEAT, INRAE, 2018. Experimental Poultry Facility, DOI: 10.15454/1.5572326250887292E12) and Sébastien Lavillatte, Maud Renouard and Edouard Guitton (PFIE, INRA, 2018. Infectiology of Farm, Model and Wild Animals Facility, https://doi.org/10.15454/1.5572352821559333E12) in charge of maintenance of the isolators and animal care. We would like to thank Irène Gabriel (INRAE Centre Val de Loire) and Marie-José Butel (University Paris Descartes) for scientific discussion and advice. We are grateful to Cindy Slugocki from the CIRM-BP Platform for checking the germ-free status.

## Supplementary files

**Supplementary figure:**
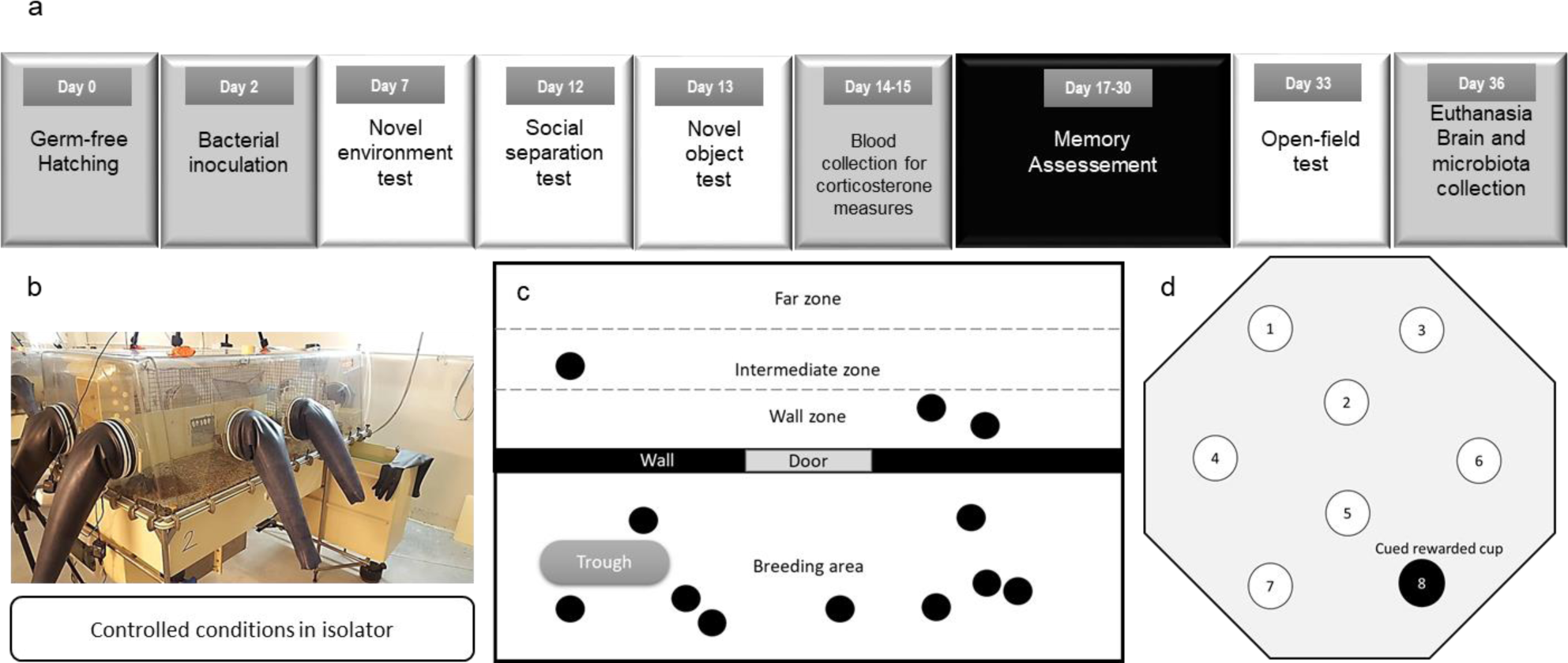
(**a**) Timing of the experimental design. Germ-free quails hatched in germ-free conditions in isolators (Day 0 = D0). After bacterial inoculation on Day 2 with cecal microbiota from a stressed or unstressed individual, the anxiety-like behavior was assessed on males and females using different behavioral tests in isolator. On Day 14 blood samples were collected for plasma corticosterone measures followed by memory tests starting on Day 17. An open-field test was performed on Day 33 to evaluate the final anxiety-like behavior level of quails before euthanasia and brains were collected for gene expression measures. (**b**) Picture of the experimental isolator used to ensure a controlled microbial environment. (**c**) The wall zone used for the novel environment test and social separation test (black circles = quails). The wall zone represents proximity with the other quails. (**d**) Schematic representation of the arena and disposition of cups used for training.

**Supplementary table 1:**
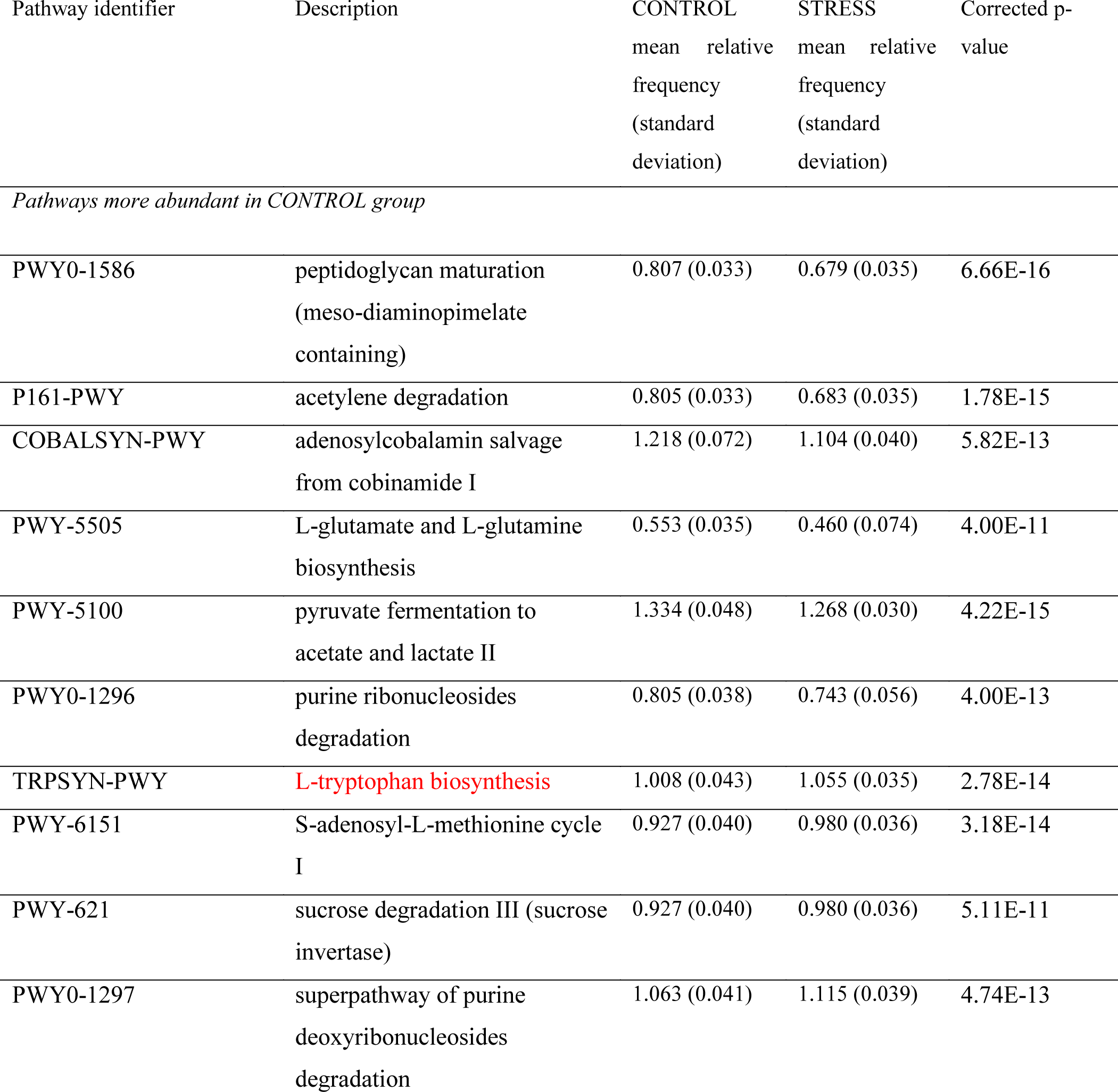
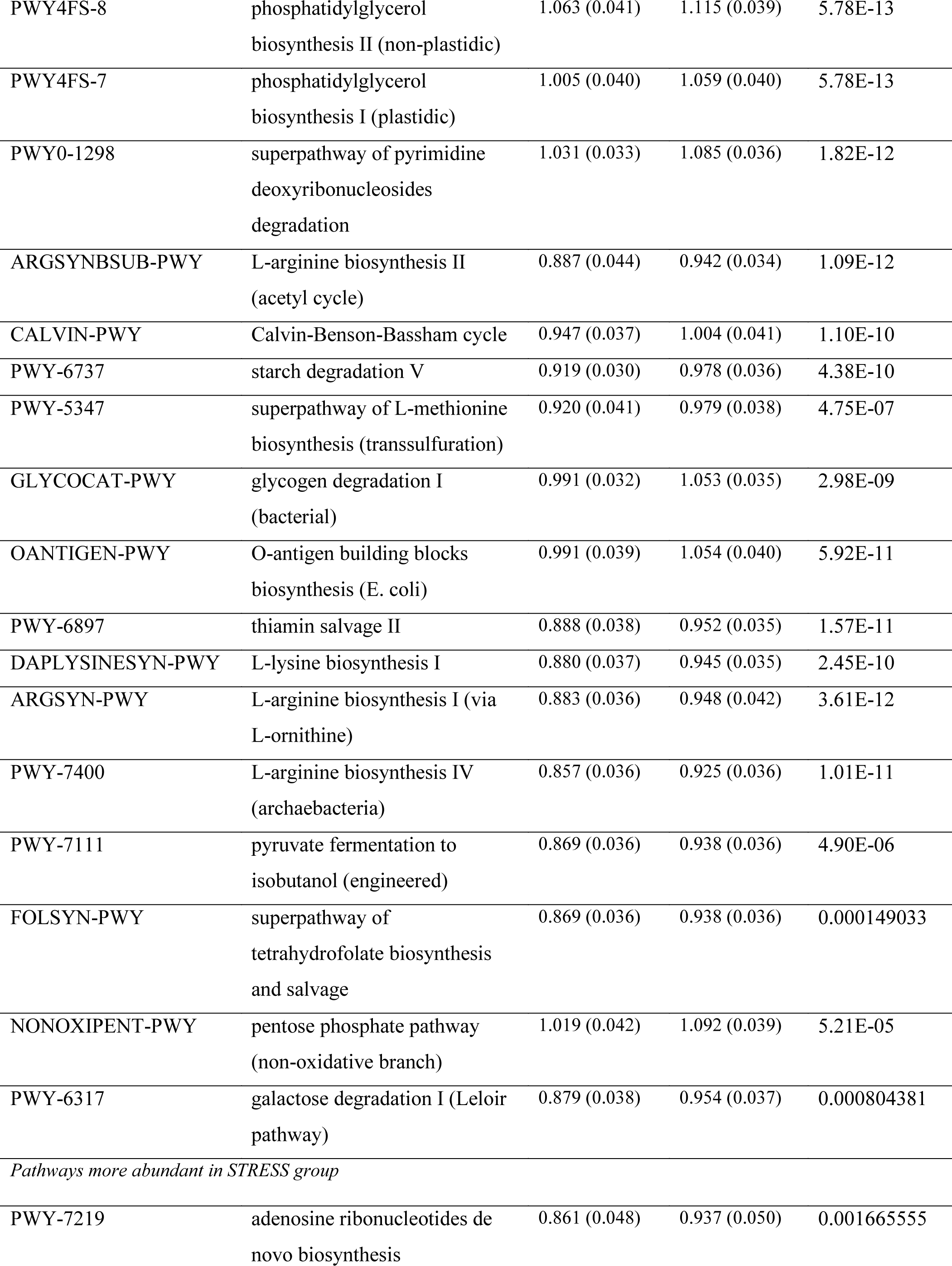
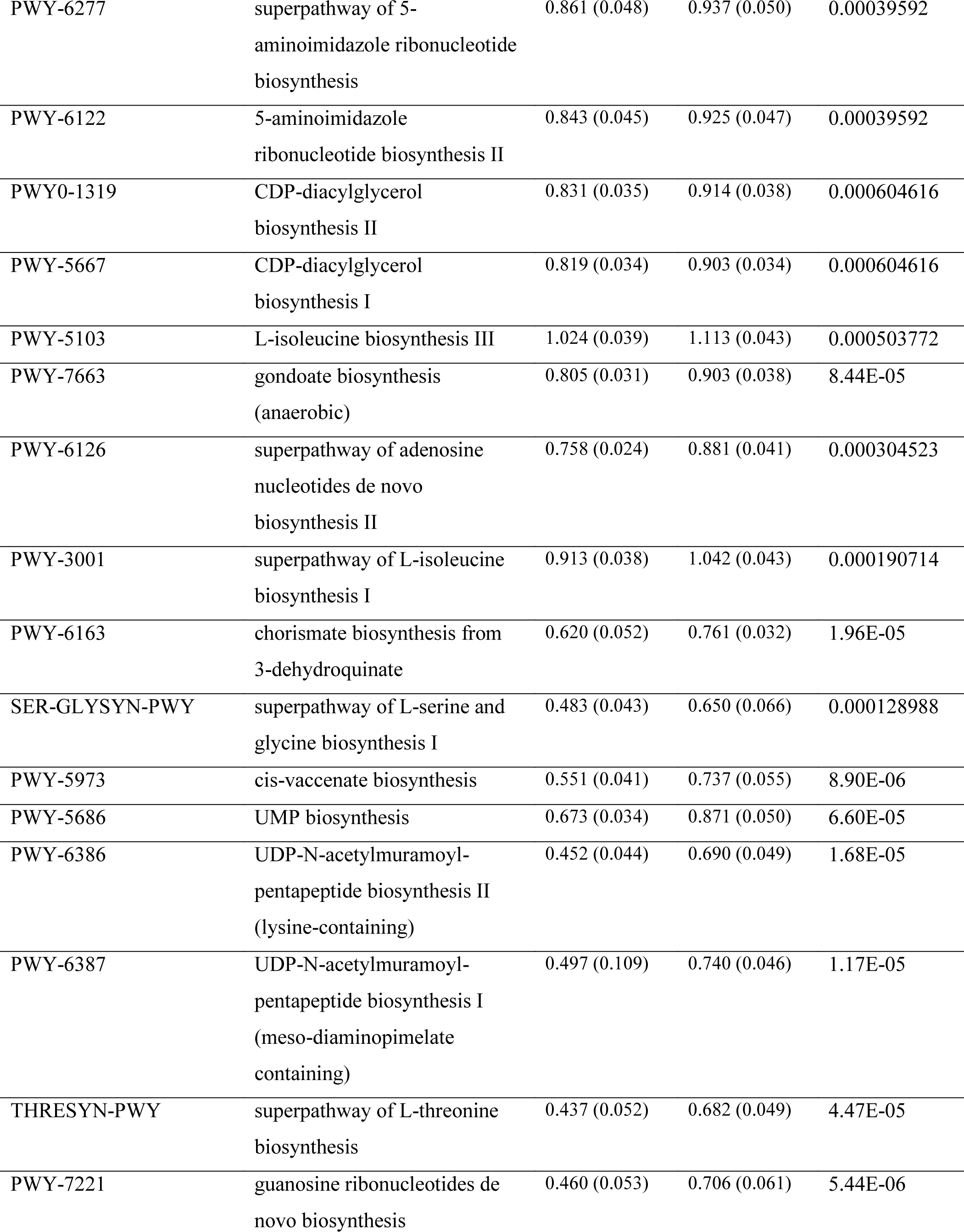
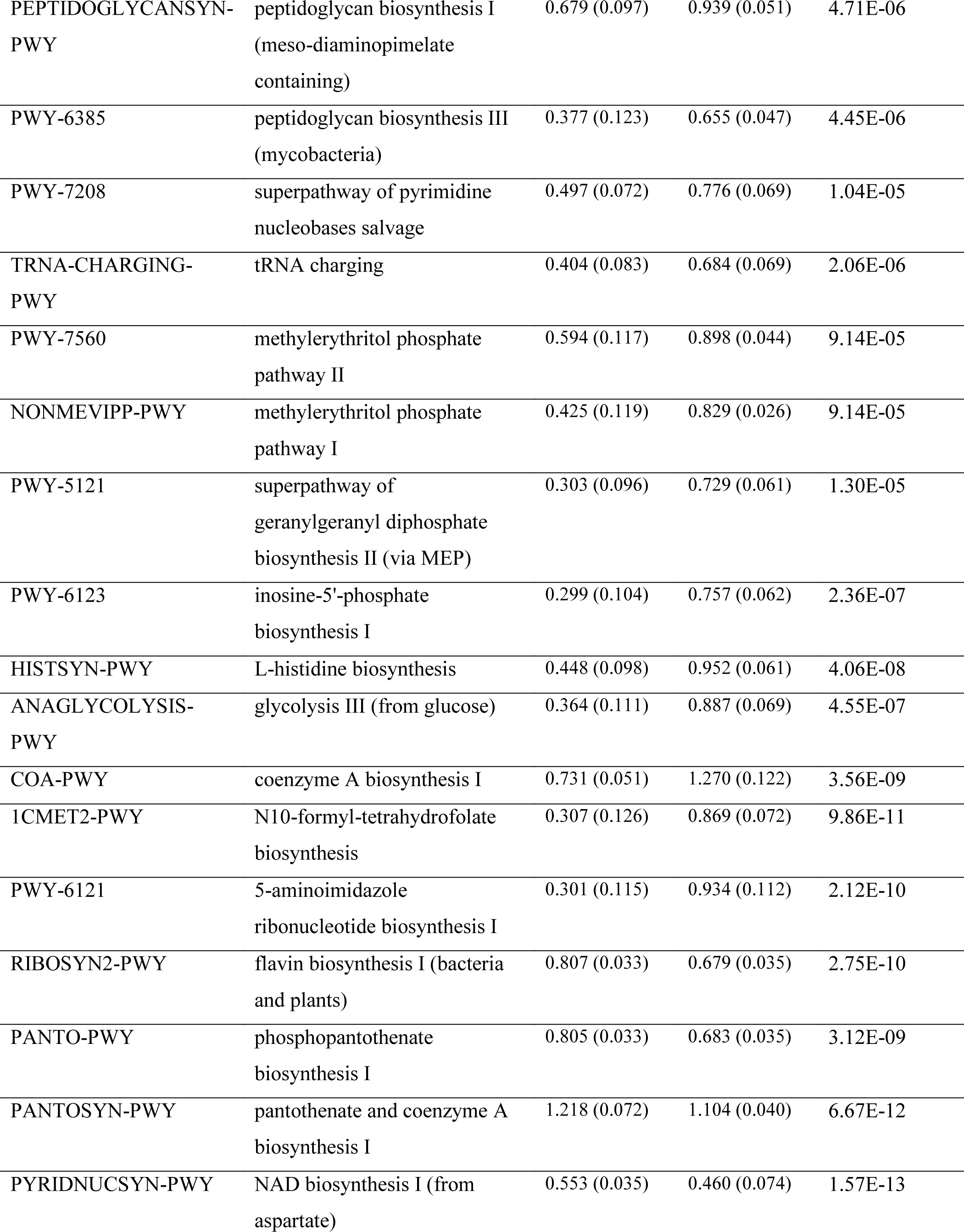
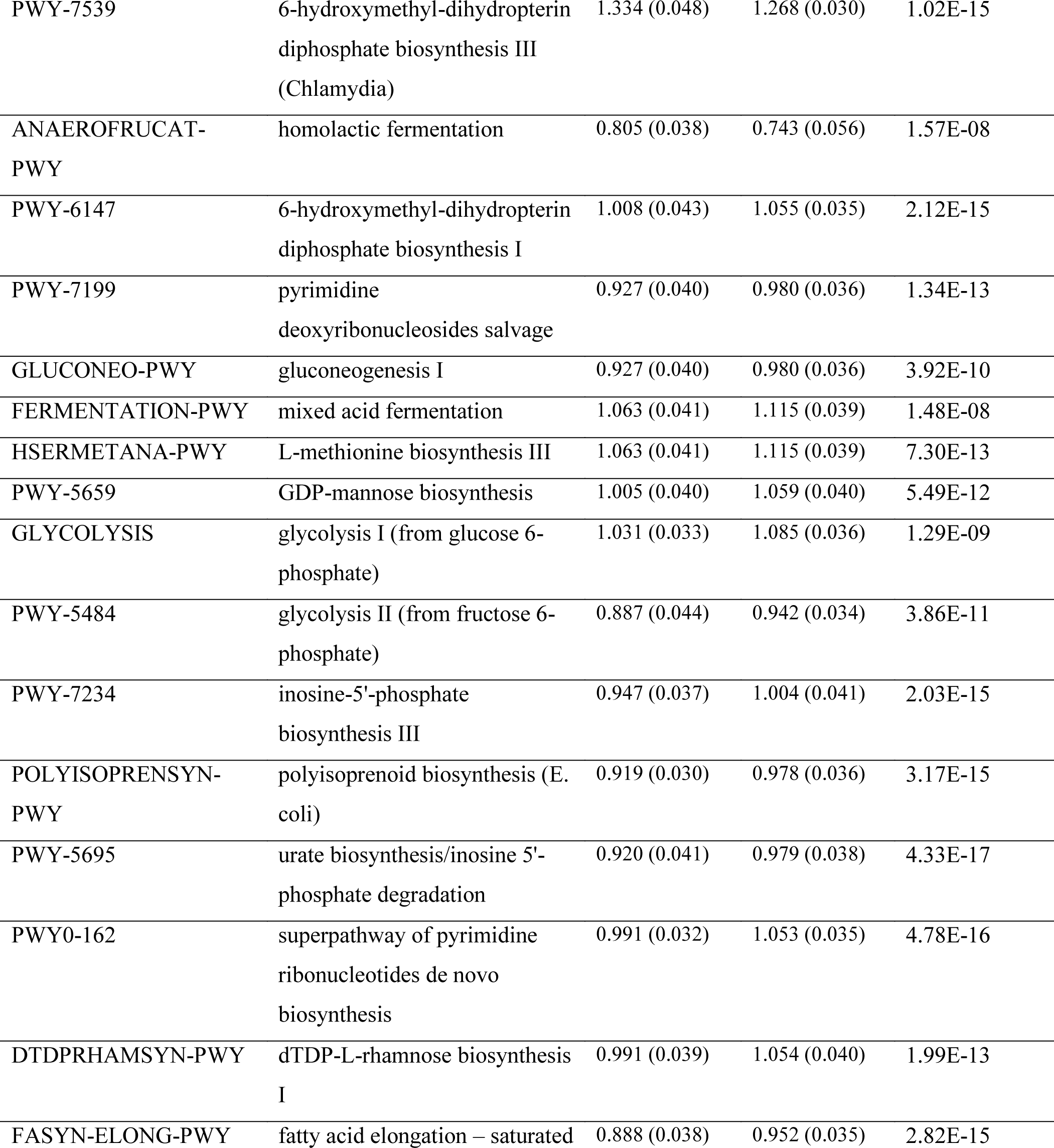
KEGG pathways predicted in the gut microbiota samples of CONTROL-T and STRESS-T groups at 14 days of age. We only report here the enriched pathways presenting significant differences between the groups (corrected *p* < 0.05) and presenting a mean relative abundance > 0.05%. The L-tryptophan biosynthesis pathway is highlighted in red.

**Supplementary table 2:** Short-chain fatty acids (SCFA) concentration in the fecal contents of quail males from the CONTROL-T (n = 15) and STRESS-T (n = 18) groups at Day 6. All data in the table are expressed as mean values ± SEM. * *p* ≤ 0.05; # *p* ≤ 0.10.

| SCFA | CONTROL-T | STRESS-T |
| --- | --- | --- |
| Acetate ($\mu\text{mol/g}$) | 4.412 ± 0.757 | 5.294 ± 0.900 |
| Propionate ($\mu\text{mol/g}$) | 0.121 ± 0.030 | 0.092 ± 0.018 |
| Butyrate ($\mu\text{mol/g}$) | 0.075 ± 0.016 | 0.078 ± 0.019 |
| Valerate ($\mu\text{mol/g}$) | 0.016 ± 0.004 | 0.005 ± 0.002 |
| Caproate ($\mu\text{mol/g}$) | 0.028 ± 0.005 | 0.016 ± 0.001 * |
| Isobutyrate ($\mu\text{mol/g}$) | 0.039 ± 0.015 | 0.021 ± 0.005 |
| Isovalerate ($\mu\text{mol/g}$) | 0.016 ± 0.004 | 0.007 ± 0.002 * |
| Isocaproate ($\mu\text{mol/g}$) | 0.015 ± 0.004 | 0.003 ± 0.001 * |
| Acetate (%) | 91.558 ± 2.014 | 94.698 ± 0.661 # |
| Propionate (%) | 2.812 ± 0.598 | 2.384 ± 0.488 |
| Butyrate (%) | 1.982 ± 0.405 | 1.589 ± 0.223 |
| Valerate (%) | 0.687 ± 0.077 | 0.236 ± 0.035 |
| Caproate (%) | 1.202 ± 0.413 | 0.519 ± 0.081 * |
| Isobutyrate (%) | 0.701 ± 0.228 | 0.521 ± 0.185 |
| Isovalerate (%) | 0.552 ± 0.201 | 0.180 ± 0.046 * |
| Isocaproate (%) | 0.505 ± 0.242 | 0.032 ± 0.020 # |

**Supplementary table 3:**
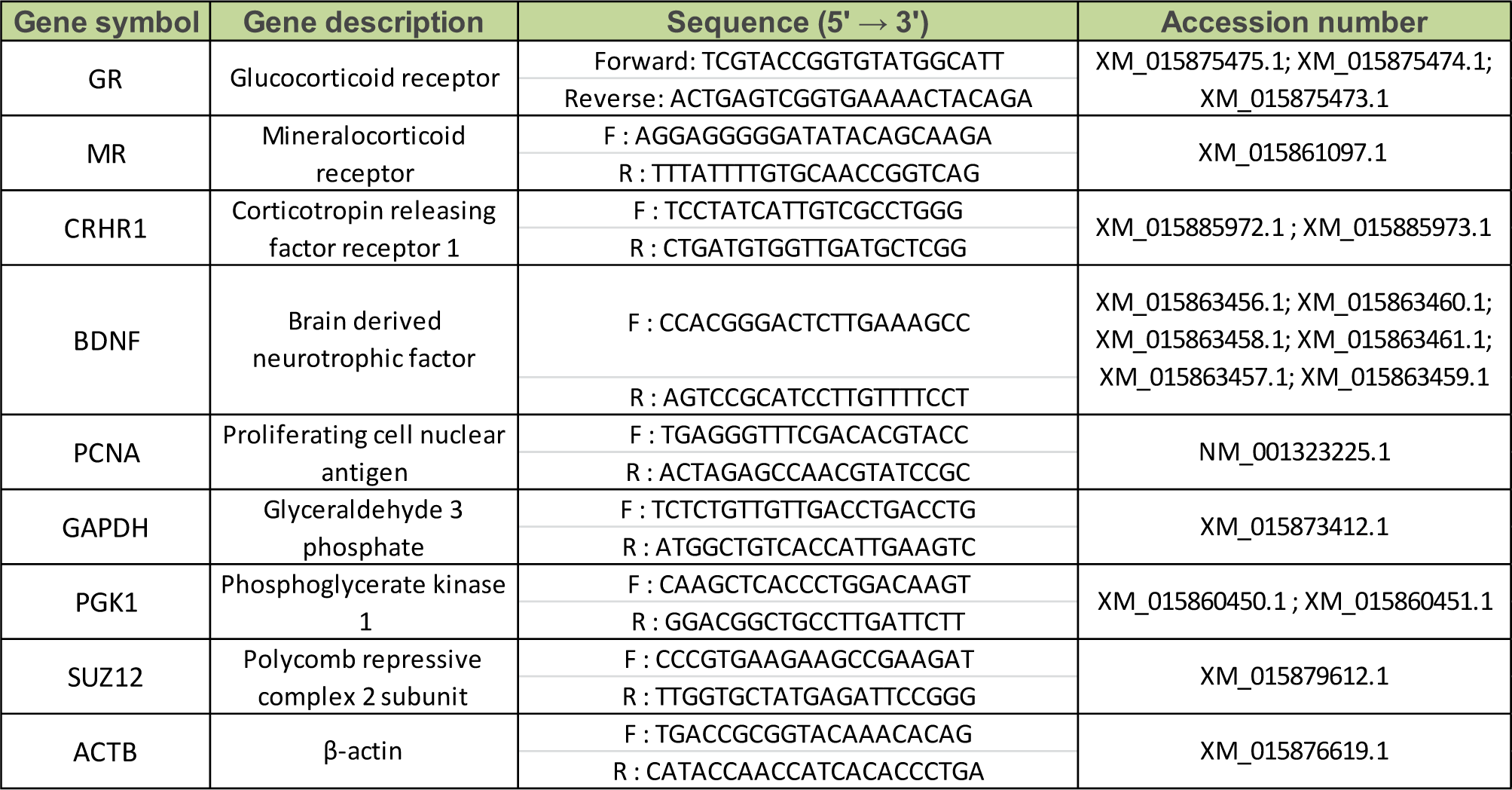
Table of sequences and number of accession of primers used for the qPCR analysis on brain.

